# Anti-cancer Drug Synergy Prediction in Understudied Tissues using Transfer Learning

**DOI:** 10.1101/2020.02.05.932657

**Authors:** Yejin Kim, Shuyu Zheng, Jing Tang, W. Jim Zheng, Zhao Li, Xiaoqian Jiang

## Abstract

**Motivation:** Exploring an exponentially increasing yet more promising space, high-throughput combinatorial drug screening has advantages in identifying cancer treatment options with higher efficacy without degradation in terms of safety. A key challenge is that accumulated number of observations in in-vitro drug responses varies greatly among different cancer types, where some tissues (such as bone and prostate) are understudied than the others. Thus, we aim to develop a drug synergy prediction model for understudied data-poor tissues as overcoming data scarcity problem.

**Results:** We collected a comprehensive set of genetic, molecular, phenotypic features for cancer cell lines from six different databases. We developed a drug synergy prediction model based on deep neural networks to integrate multi-modal input and utilize transfer learning from data-rich tissues to data-poor tissues. We showed improved accuracy in predicting drug synergy in understudied tissues without enough drug combination screening data nor after-treatment transcriptome. Our synergy prediction model can be used to rank synergistic drug combinations in understudied tissues and thus help prioritizing future in-vitro experiments.

**Availability and Implementation:** Our algorithm will be publicly available via https://github.com/yejinjkim/drug-synergy-prediction

## Introduction

Drug discovery is an expensive and time-consuming process. It takes many years and costs billions of dollars for a drug to be approved. Systematic integration of previous results and knowledge might change the game by identifying highly promising drugs and their combinations to save cost and speedup discovery (e.g., repositioning FDA approved drug lists might reduce the risk of drug toxicity). High-throughput drug screening using computational approaches has the potential to substantially improve cost-efficiency by automatically estimating drug sensitivity based on genomic and pharmacological data.^1–7^ These computational methods utilize drug sensitivity data at certain cell lines and predict unknown drugs that potentially have high sensitivity in other cell lines. In addition to the drug-cell line interaction, another important drug response type is *synergy* between drugs. High-throughput combinatorial drug discoveries throw light on treating complicated multifactorial diseases, including diabetes and metabolic syndrome, cardiovascular disease, cancer, and neurodegenerative disorders.^8–15^ Through exploring an exponentially larger, more challenging, yet promising space, the drug combination strategies have advantages in identifying candidates with higher efficacy without degradation in terms of safety.^10,16–18^ They show great promises in finding potent drug combinations that cannot be identified by one drug - one target approaches. For example, certain high efficacy drug combination can be composed of two or more drugs that each does not have a significant treatment effect (e.g., synergistic drug interactions for treating cancer^19^). The result can be attributed to additive or synergistic drug combinations. The synergistic ones, which offer stronger combined effects than simply adding up effects of individual drugs, are of particular interest and associated with: (1) complementary actions when two or more drugs target multiple points along the same protein or pathway, (2) anti-counteractive actions when a secondary drug targets a biological response to the primary drug, and (3) facilitating actions are combinations where a secondary drug promotes the activity of the primary therapeutic.^8^

A key challenge in computational drug synergy prediction model is that accumulated number of observations in in-vitro drug responses varies greatly for different tissues. Traditional methods target at studying commonly observed tissues such as breast, kidney, skin, lungs.^8–15^ These methods take known drug response data at certain cell lines and attempt to find other drugs responses at other cell lines within the data-rich tissues. The understudied tissues include bone, prostate, and pancreas (Fig. S1a and S1b). A number of obstacles impede the drug response study in these tissues. For example, bone cancer drugs response has been poorly understood due to physical difficulty of culturing bone tissues as cell lines, the rarity of the tumors in sarcoma type, the difficulties of obtaining tumor tissue fragments from human patients for bone metastasis, and thus the limited number of cell models,^20^ despite the fact that the most common form of bone cancer (i.e., bone sarcomas and bone metastasis) are considered deadly and incurable. For prostate cancer, the difficulty of obtaining its cell lines has been due to heterogeneity in prostate gland’s cell and hormone responsiveness.^21,22^ The lack of cell line models makes the high-throughput screening difficult, which in turn makes these tissues even more understudied. There is, therefore, a critical need to develop a computational drug combination prediction tool for the understudied tissues.

Our ultimate goal is to rank drug combinations that are highly likely to be synergistic in understudied tissues in silico and thus help prioritizing future in-vitro experiments. To attain this goal, we aimed to develop a drug response prediction model for the understudied tissues. The most critical challenge in developing the model is data scarcity. Understudied tissues inevitably lack enough training data; they have a limited number of experimental observations and also lack of important features such as post-treatment transcriptome. The post-treatment transcriptome has been a popular tool for drug repositioning as it directly quantifies gene expression levels and measures biological effects of a compound.^23^ A potential way to mitigate this data scarcity problem is to utilize information from the data-rich tissues to the data-poor tissues as these different tissues share biological commonality partly in terms of gene expression and therefore respond drugs in similar ways.^24^ Several previous studies support that anti-cancer drug sensitivity in cell lines is not tissue-specific and that tissue-specific drugs can bring additional benefits to other tissues. ^25,26^ Accordingly we proposed to utilize interaction between drugs and cell lines learned from the data-rich tissues to help increase the performance of the data-poor tissue using *transfer learning*. Transfer learning is to transfer knowledge (in the form of parameters in machine learning model) from previously learned model (teacher model) with large data to new model (student model) with less data. Main obstacle in applying the transfer learning to our problem is to learn “transferable” knowledge in data-rich tissues in spite of the differences in underlying biological mechanism between data-rich and data-poor tissues. To learn generalizable and thus transferable knowledge on drug target and cell line gene expressions in the teacher model, we integrated all types of available tissues in one model with a large dataset (i.e., 4,150 drugs; 112 cell lines; 16,816 monotherapy sensitivity; 1,163,498 combinations synergy) so that the teacher model can simulate non-tissue-specific drug-cell interaction. To supplement the lack of post-treatment transcriptome, we maximally utilized various types of multi-modal inputs (molecular, genetic, phenotypic features) and multiple outputs (drug sensitivity and synergy) by harmonizing six different databases. For prediction model we used deep neural networks to incorporate the different types of multi-modal inputs and multi-task learning to jointly incorporate the two responses (monotherapy sensitivity and combination synergy) at the same time.

## Results

### Prediction accuracy

We evaluated the accuracy of our drug combination prediction models. We measured accuracy of predicted monotherapy sensitivity and combination synergy. We performed the experiments with regression and classification task. Accuracy measures were mean squared error (MSE) for regression task that predicts sensitivity and synergy values; and area under the receiver operating curve (AUC) and normalized discounted cumulative gain (NDCG) as a ranking performance for classification task that predicts or classifies binarized sensitivity and synergy. We set binarizing threshold at 10 following the previous work.^27^ We compared the accuracy as adding more input features: ID, ID+F, and ID+F+G, where ID models have inputs of {drug ID} and {cell line ID}; ID+F models have inputs of {drug ID, MACCS fingerprints, SMILES NLP features} and {cell line ID, tissue, cancer types}; ID+F+G models have inputs of {drug ID, MACCS fingerprints, SMILES NLP features, target genes} and {cell line ID, tissue, cancer types, gene expression profiles}. We compared our accuracy with baseline method - DeepSynergy that utilizes deep neural networks using drug’s descriptors and cell line’s gene expression.^10^ Benchmark accuracy values of various machine learning methods (including support vector machines, gradient boosting machines, and random forest) are listed in this paper.^10^

First experiment was to evaluate the accuracy of models trained and tested with data-rich tissues (Table 1). We achieved {0.95 AUC, 1.0 NDCG, 115 MSE} for sensitivity and {0.89 AUC, 0.84 NDCG, 178 MSE} for synergy. We observed that adding multimodal features increase accuracy by comparing accuracy between ID, ID+F, and ID+F+G. Compared to the baseline method DeepSynergy,^10^ our model achieved significantly lower MSE and similar AUC, even though our classification task was difficult as setting binarized threshold lower-- (our model=10 vs DeepSynergy=30), thanks to 20-times larger scale datasets and more diverse multi-modal features (such as target genes and native SMILES).

**Table 1.** Drug response prediction accuracy for data-rich tissues. Trained with data-rich tissues and tested with data-rich tissues. For classification task, AUC and NDCG were computed after binarizing sensitivity and synergy at threshold 10. AUC=area under the receiver operating curve; NDCG=normalized discounted cumulative gain; MSE=mean squared error.

| Different input features | Monotherapy Sensitivity |  |  | Combination Synergy |  |  |
| --- | --- | --- | --- | --- | --- | --- |
|  | AUC | NDCG | MSE | AUC | NDCG | MSE |
| ID | 0.9302 | 1.0000 | 137.6218 | 0.8570 | 0.8132 | 184.8153 |
| ID+F | 0.9423 | 1.0000 | 119.6132 | **0.8861** | **0.8377** | 177.8911 |
| ID+F+G | **0.9498** | 1.0000 | **115.3092** | 0.8844 | 0.7332 | **174.3452** |

In next experiments we moved our focus to understudied tissues. We transferred the model parameters from data-rich tissues to bone and prostate respectively. We compared accuracy with and without the transferred model parameters. We found that transfer learning increases accuracy in most cases in both bone and prostate (Table 2). In bone, we achieved 0.94 AUC for sensitivity. For synergy, we achieved 0.83 AUC from ID+F+G model after transfer learning but the accuracy without transfer learning was also comparable. In prostate tissue, we achieved 0.96 AUC for sensitivity and 0.86 AUC for synergy using ID+F+G model after transfer learning.

**Table 2.** Drug response prediction accuracy for bone and prostate cancer.

|  | Different input features | Monotherapy Sensitivity |  |  | Combination Synergy |  |  |
| --- | --- | --- | --- | --- | --- | --- | --- |
|  |  | AUC | NDCG | MSE | AUC | NDCG | MSE |
| Bone | Without transfer learning (trained and tested with bone) |  |  |  |  |  |  |
|  | ID | 0.9319 | 0.9636 | 278.9249 | 0.8096 | 0.4215 | 146.1300 |
|  | ID+F | 0.9285 | 1.0000 | 178.8830 | 0.8057 | 0.3966 | **78.1472** |
|  | ID+F+G | 0.9273 | 1.0000 | 221.1020 | 0.7906 | **0.4939** | 103.0315 |
|  | With transfer learning (trained with data-rich tissues, retrained with bone, and tested with bone) |  |  |  |  |  |  |
|  | ID | 0.9325 | 1.0000 | 283.1641 | **0.8564** | 0.3609 | 149.1630 |
|  | ID+F | **0.9363** | 0.9290 | **164.8712** | 0.7974 | 0.3667 | 84.7019 |
|  | ID+F+G | 0.9295 | 1.0000 | 203.6013 | 0.7921 | 0.3599 | 89.5048 |
| Prostate | Without transfer learning (trained and tested with prostate) |  |  |  |  |  |  |
|  | ID | 0.9477 | 1.0000 | 101.5557 | 0.7893 | 0.4671 | 212.3478 |
|  | ID+F | **0.9615** | 1.0000 | 80.7862 | 0.8120 | 0.3492 | 176.1962 |
|  | ID+F+G | 0.9591 | 1.0000 | 92.2249 | 0.8375 | 0.2921 | 172.9621 |
|  | With transfer learning (trained with data-rich tissues, retrained with prostate, and tested with prostate) |  |  |  |  |  |  |
|  | ID | 0.9544 | 1.0000 | 91.1933 | 0.7977 | 0.4354 | 205.7588 |
|  | ID+F | 0.9596 | 1.0000 | **70.4784** | 0.8295 | **0.5075** | **162.8379** |
|  | ID+F+G | 0.9596 | 1.0000 | 75.0662 | **0.8597** | 0.4710 | 164.1078 |

### Ranking

As a case study, we listed predicted drug combinations for bone using ID+F+G model. We selected top 20 combinations based on estimated probability of being synergistic (Table S4). Among the top 20 five combinations were actually synergistic (5/20=25% hit). Considering the fact that only 1.56% of combinations are synergistic in bone, our model effectively ranked synergistic drug combinations. We also listed predicted drug combinations for prostate using ID+F model. We selected top 20 combinations based on estimated probability of being synergistic (Table S4). Among the top 20 ten combinations were actually synergistic (10/20=50% hit). In prostate tissue, only 2.15% of combinations are synergistic.

## Discussion

The objective of this study was to develop the drug combination response prediction model, which can be particularly used even in the understudied tissue with less observation. To meet this end, we i) collected a comprehensive set of genetic data from multiple databases, ii) integrated multi-modal source of features using deep neural networks, iii) transferred the prediction model from data-rich tissues to data-poor one with fine tuning. As a result, the proposed model predicted drug response accurately with 0.95 AUC for sensitivity and 0.89 AUC for synergy in data-rich tissues. For the data-poor tissues, we achieved 0.94 AUC for sensitivity and 0.83 AUC for synergy in bone; 0.96 AUC for sensitivity and 0.86 AUC for synergy in prostate using transfer learning. When compared accuracy with and without transfer learning, transfer learning consistently increased accuracy in prostate but not in bone. We hypothesized that drugs and bone cancer cell lines interact in different way as bone is physically different from other tissues.

Main contribution of our study is that we tackle understudied but critical tissues for drug response prediction. The difficulty of obtaining cell lines in these understudied tissues has limited the in-vitro experiments, which consequently become an obstacle to drug development in these tissues. With the help of computational drug response model, researchers can effectively prioritize drug combinations that are highly likely to be synergistic. As we focused on understudied tissues, we should avoid using post-treatment gene expression profiles as an input feature. The post-treatment gene expression is the most powerful features to estimate drug response, but it is only accessible after drug compounds are tested in cell line.^9^ In understudied tissues, it is even more difficult to access the post-treatment results. Our drug response prediction model was able to successfully bypass the post-treatment features while achieving competitive accuracy.

Although our study’s focus is on predicting synergy in understudied tissues, our model achieved improved accuracy even on general data-rich tissues than that of baseline model. This increased accuracy is possibly due to the large-scale and multi-modal data. We incorporated diverse and comprehensive multi-modal source of features (e.g., SMILES, drug’s target genes, cell line’s gene expressions) by integrating several publicly available databases, and used a total of 23 synergy databases (whereas the baseline study used only one database). This large-scale data collection allowed us to maximize the power of data-driven computational model based on deep neural networks. In addition, we used state-of-the-art NLP encoder to incorporate drug’s native chemical features. We also carefully designed the model structure to avoid redundant parameters, whereas the baseline method set redundant parameters for each drug in pairs (e.g., DeepSynergy model differentiates drug ordering).^10^

Our model was designed to predict monotherapy sensitivity and combination synergy simultaneously. Here monotherapy sensitivity was an auxiliary output to boost accuracy of synergy prediction. A previous study uses monotherapy sensitivity as an input feature to predict synergy,^9^ but understudied tissues do not have enough experimental observation including monotherapy response features. To utilize this partial input feature in data-rich tissues (not in data-poor tissues), we used the monotherapy sensitivity as an auxiliary output in data-rich tissue model so that we can still transfer the information to data-poor tissues via the model parameters when minor tissues do not have monotherapy sensitivity.

In conclusion, our model is an end-to-end drug synergy prediction model that learns interaction between drugs (in terms of past drug history, molecular structure, and target genes) and cell line (in terms of past cell line history, tissue, cancer type, and baseline gene expression). Based on the fact that different tissues share common gene expression and therefore respond drugs in similar ways, we used transfer learning from data-rich tissues to data-poor tissues to make the synergy prediction model work in data-poor tissues. As a future work, our drug prediction model for understudied tissues can potentially shed light onto other diseases that share drug targets and underlying mechanism and offer a novel way of efficient and low-cost drug discovery.

## Materials

### Drug sensitivity and synergy

We used a publicly available large-scale drug synergy database from DrugComb Portal,^28^ which combines 23 drug synergy studies into one integrated database. The number of unique drugs and cell lines were 4,150 and 112, respectively (Fig. S1). There are 2,843 experimental drugs and 1,307 FDA-approved drugs. There were a total of 710,242 monotherapy sensitivity (a pair of drug and cell line) and 466,259 drug combinations synergy (a triplet of drug1, drug2, and cell line). For monotherapy sensitivity we calculated relative inhibition (RI) from dosage combination matrix (Supplementary 1). For drug combinations synergy we used Loewe synergy score (Fig. S1), which ranged from −100 (antagonistic effect) to 75 (strong synergistic effect).^29^ Loewe synergy score is to quantifies the excess over the expected response if the two drugs are the same compound.^30,31^ We selected Loewe score for comparison with the baseline study.^10^

### Drug’s features

We extracted drug’s molecular and genomic features. For molecular features, we used both Molecular ACCess System (MACCS) fingerprints^32^ and native chemical compounds. MACCS fingerprints contains 166 chemical structures such as the number of oxygens, S-S bonds, ring. We used RDKit (http://www.rdkit.org) to extract MACCS fingerprints. In addition, we represented drug as a native chemical structure using SMILES. SMILES is a linear notation to represent chemical compound in a unique way; in the SMILES representation atoms are represented as their atomic symbols (e.g., c for carbon); special characters are also used to represent relationship (e.g., “=”: double bonds; “#”:triple bonds; “.”:ionic bond; “:”: aromatic bond)^33^. SMILES can provide richer features space that strictly represent functional substructures and express structural differences such as compound’s chirality ^34^. We used the state-of-the-art natural language processing model, Transformer, to encode the SMILES sequence.^35^ For genomic features, we integrated three different drug databases - DrugBank,^36^ Therapeutic Targets Database (TTD),^37^ and NIH-LINC^38^ - for complete view of drug’s target genes (Table S1). We filtered out non-human target genes in TTD.

### Cell line’s features

We used cell line’s tissue/cancer type and genomic features. There were 14 tissues including lung, ovary, and skin and 14 cancer types including carcinoma, adenocarcinoma, and melanoma (Table S2). We also extracted gene expression profiles by Fragments Per Kilobase of transcript per Million reads mapped (FPKM) from CCLE and COSMIC.^39,40^ We found 75 cell lines with 37,260 genes from Broad Institute and 25 cell lines with 35,004 genes from Sangar (Table S1). In total, we found 88 cell lines with 22,586 genes after excluding zero-variance genes. We normalized the FPKM values into z-score in a gene-wise manner as FPKM varies greatly depending on gene. We only used baseline (before-treatment) gene expression profiles without after-treatment gene profiles because our objective is to test our models in understudied tissues, which rarely has after-treatment gene profiles.

## Methods

### Method overview

Our objective is to predict whether unobserved drugs and their combination has sensitivity and synergistic effects in a given cell line and provide a list of combinations that can researchers prioritize for experiments. We formulated it as a prediction problem. Given an experimental block *x*_*n*_ : = {*d*_*i*_, *d*_*j*_, *c*_*k*_}of drug combination (*d*_*i*_ , *d*_*j*_) and cell line *c*_*k*_, the prediction model is a function *f* such that

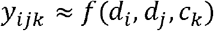

where *Y*_*ijk*_ is synergy score of the drug combination. For monotherapy sensitivity in the cell line, the prediction model is *g*:

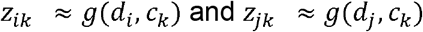

where *z*_*ik*_ is the sensitivity score of drug *d*_*i*_ in cell line*c*_*k*_.We jointly optimize sensitivity and synergy as multi-task learning. To incorporate multimodal source of inputs and formulate nonlinear relationship between chemical and genomic features, we used deep neural networks as the prediction function. Our prediction model consists of drug encoders (Fig. 1a), cell line encoder (Fig. 1b), and merging layers (Fig. 1c) for final prediction in an end-to-end manner. Using the estimated drug response, we can rank drug combination at a cell line that are expected to have synergistic effect.

**Figure 1.**
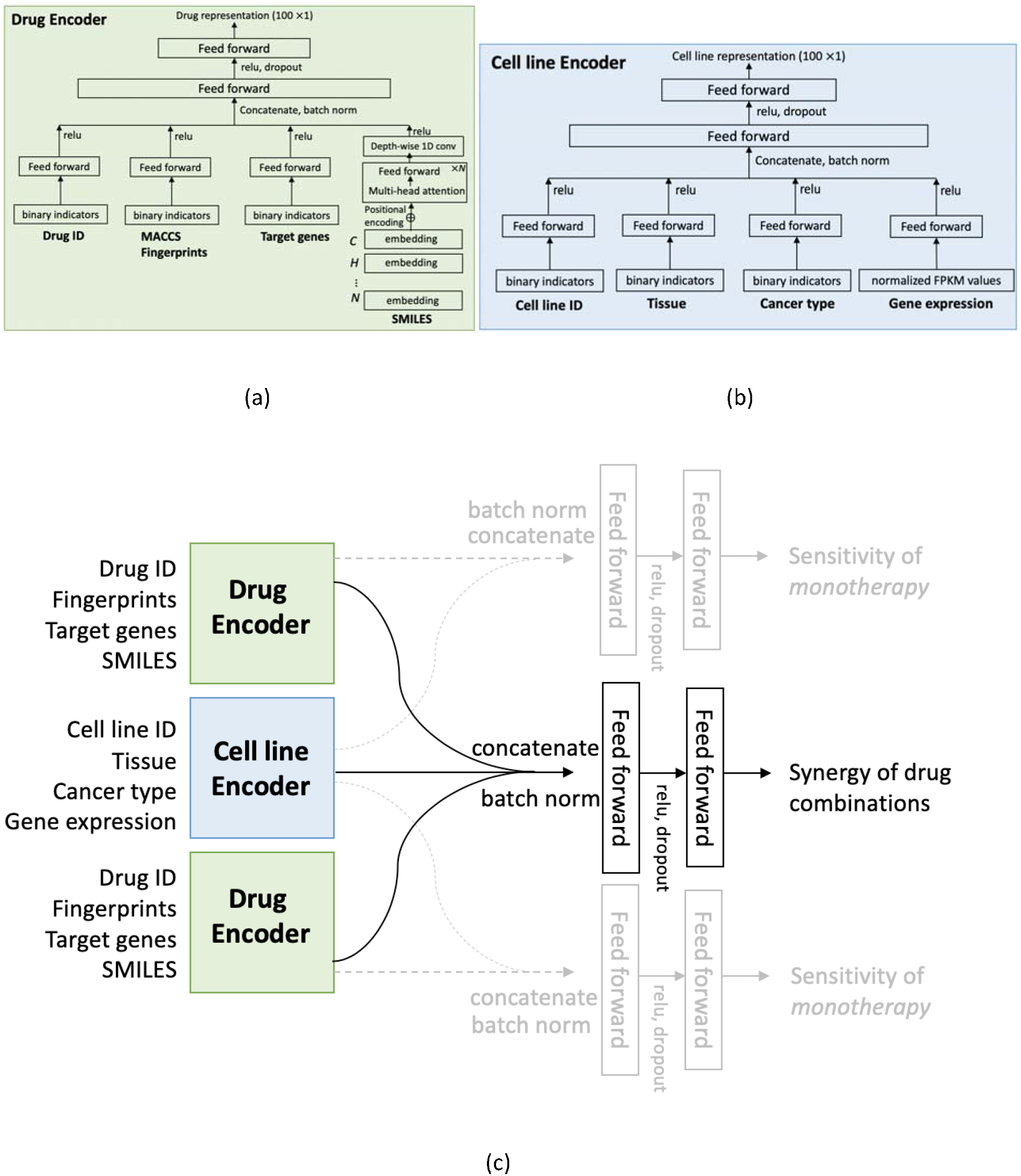
Drug synergy prediction model. (a) Drug encoder. It learns an embedding representation of a drug. Inputs are drug ID, MACCS fingerprints, canonical SMILES, and target genes. (b) Cell line encoder. It learns an embedding representation of cell line. Inputs are cell line ID, tissue, cancer type, and gene expressions. (c) Merging drug encoder and cell line encoder. Sensitivity was an auxiliary output to boost synergy prediction.

### Drug encoder

Drug encoder learns an embedding representation of each drug *d*_*i*_ or *d*_*j*_. Each drug’s features are {drug ID, MACCS fingerprints, canonical SMILES, target genes} (Table S3). One-hot vector of drug ID was represented as the same sized embedding. Binary indicator of MACCS fingerprints were used as raw input. Binary vector of target genes was represented as half-sized embedding via linear transformations (Fig. 1a).

As SMILES has variable length, we need more careful consideration. We used Transformer encoder, a natural language processing model that convert the sequence into a representation.^35^ Each symbol in SMILES were first represented as one-hot vectors with size of (#SMILES length * #unique symbols), where #unique symbols was 48 and the maximum SMILES length was 288. The symbol’s one-hot vector was then represented as embedding. This embedding sequence was fed into a separate Transformer encoder, which consists of multi-head attention layer and feed-forward layer with repeated residual connections. Once we derived all the embedding representations, they were concatenated into one and fed into two feed forward layers with Relu activation and dropout.

### Cell line encoder

Cell line encoder learns an embedding representation of each cell line *c*_*k*_. Each cell line’s features are {cell line ID, tissue, cancer type, gene expressions} (Table S3). To combine the multimodal inputs, we derived embedding from each source and merged into one embedding. One-hot vector of cell line ID, tissue, and cancer types were represented as the same sized embedding, respectively. The gene expression of each cell line was represented as normalized FPKM values, which were fed into feed forward layer with Relu activation and dropout. We concatenated all four embeddings into one and fed them into two feed forward layers with Relu activation and dropout (Fig. 1b).

### Merging layers

For synergy prediction, we merged embedding representations of into one and fed them into two feed forward layers with Relu activation. For sensitivity prediction, we merged embedding representation from ( ) and ( ), respectively, and fed them into two feed forward layers (Fig. 1c).

### Training loss

We trained the model as multi-task learning that predicts synergy and sensitivity simultaneously. For combination synergy prediction, the training loss was mean squared error (MSE):

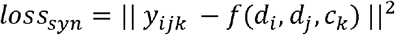

where *Y*_*ijk*_ is the synergy score. The training loss for monotherapy (*d*_*i*_ , *c*_*k*_) and (*d*_*j*_,*c*_*k*_) was similarly defined:

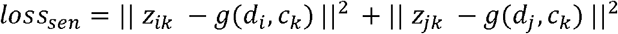

for all drugs and cell lines in training set. In addition to these regressions, we also developed classification model. We first binarized the drug responses by setting threshold. That is, 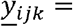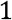 if *Y*_*ijk*_ > *threshold*; 0 otherwise, and 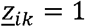 if *z*_*ik*_ > *threshold*; 0 otherwise. The threshold was set at 10 following the previous study.^27^ Then the classification model’s training loss was binary cross entropy:

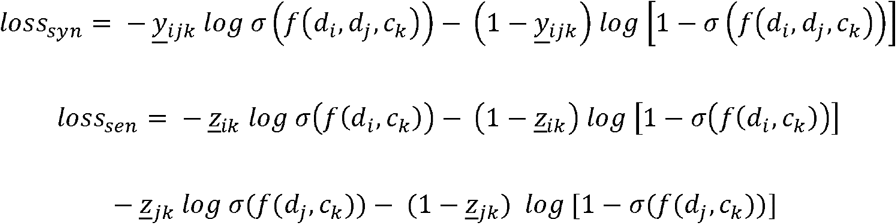

where *σ* is sigmoid function. We alternated the two optimization tasks with respect to synergy or sensitivity in every epoch. That is, we first minimized *loss*_*syn*_with all training batches and then switched to *loss*_*sen*_ for the next training round. Optimizer was adaptive gradient descent with autograd in Pytorch 1.3.0.

### Transfer learning

We focused on transfer learning from data-rich tissues to data-poor tissues with less observation: bone and prostate (Fig. 2a). We first derived general drug response prediction model with other data-rich tissues to learn the underlying mechanism between drugs and cell line (using training set in data-rich tissues), then transferred the model parameters into the understudied tissues. We retrained the model with respect to training set for the respective understudied tissue and evaluated the model with the test set in each tissue (Fig. 2b). Note that, in language modeling and image processing, this transfer learning and fine-tuning techniques has improved the classification accuracy of tasks with few observations to train models.^41^

**Figure 2.**
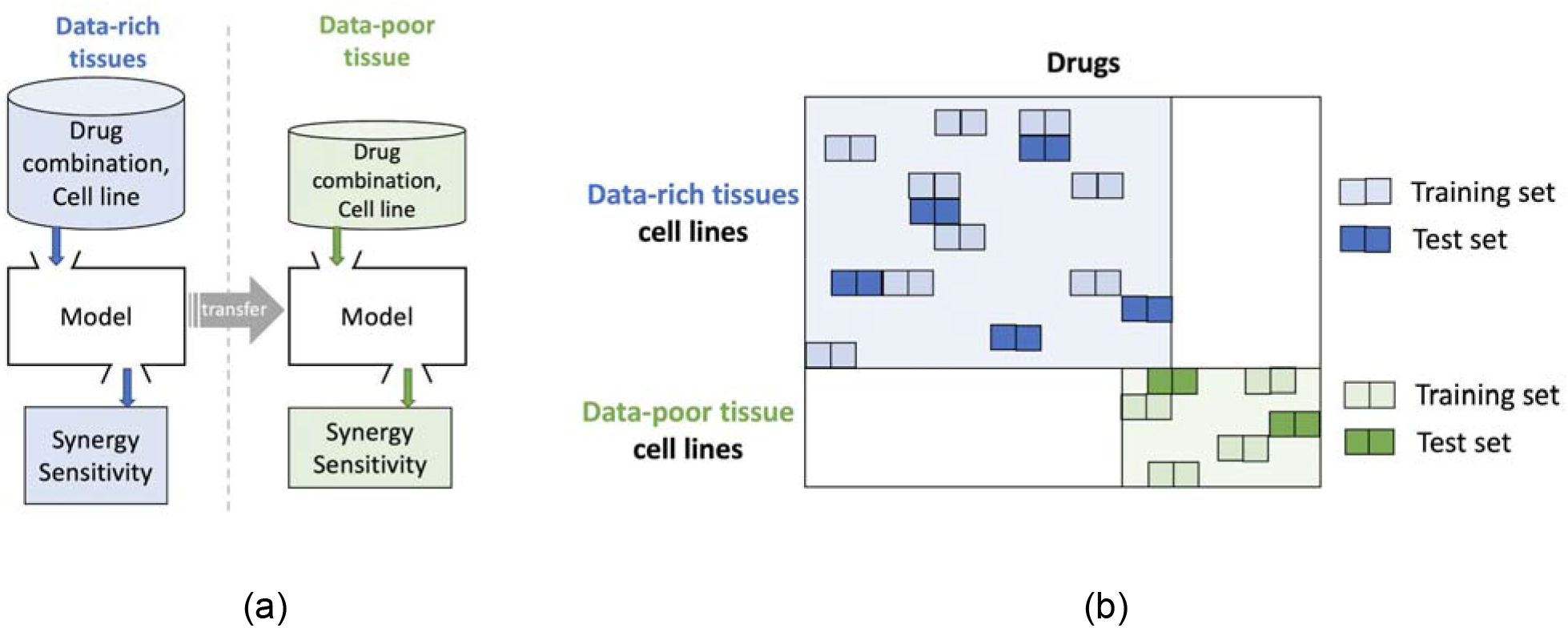
Transfer learning for understudied tissue (with an example of bone tissue). (a) Transfer model parameters from data-rich tissues to data-poor tissue such as bone and prostate (b) Train/test split for data-rich tissues and data-poor tissue. Cell lines from different tissues are disjoint. Some cell lines from different tissues sometimes share drugs (e.g., 1,437 drugs for bone and 127 drugs for prostate).

### Evaluation

We evaluated the fine-tuned prediction model with randomly selected 20% test set in each understudied tissue. To measure the prediction accuracy, we reported MSE for regression task. For classification task, we measured (pointwise) accuracy using area under the receiver operating curve (AUC). In addition, we measured ranking performance using normalized discounted cumulative gain (NDCG, Supplementary 2)

